# Differential Patterns of Synaptic Plasticity in the Nucleus Accumbens Caused by Continuous and Interrupted Morphine Exposure

**DOI:** 10.1101/2022.11.01.514765

**Authors:** Emilia M. Lefevre, Elysia A. Gauthier, Lauren L. Bystrom, Jordan Scheunemann, Patrick E. Rothwell

**Affiliations:** Department of Neuroscience, University of Minnesota, Minneapolis, MN

## Abstract

Opioid exposure and withdrawal both cause adaptations in brain circuits that may contribute to abuse liability. These adaptations vary in magnitude and direction following different patterns of opioid exposure, but few studies have systematically manipulated the pattern of opioid administration while measuring neurobiological impact. In this study, we compared cellular and synaptic adaptations in the nucleus accumbens shell caused by morphine exposure that was either continuous, or interrupted by daily bouts of naloxone-precipitated withdrawal. At the behavioral level, continuous morphine administration caused psychomotor tolerance, which was reversed when the continuity of morphine action was interrupted by naloxone-precipitated withdrawal. Using *ex vivo* slice electrophysiology in female and male mice, we investigated how these patterns of morphine administration altered intrinsic excitability and synaptic plasticity of medium spiny neurons (MSNs) expressing the D1 or D2 dopamine receptor. We found that morphine-evoked adaptations at excitatory synapses were predominately conserved between patterns of administration, but there were divergent effects on inhibitory synapses and the subsequent balance between excitatory and inhibitory synaptic input. Overall, our data suggest that continuous morphine administration produces adaptations that dampen the output of D1-MSNs, which are canonically thought to promote reward-related behaviors. Interruption of otherwise continuous morphine exposure does not dampen D1-MSN functional output to the same extent, which may enhance behavioral responses to subsequent opioid exposure. Our findings support the hypothesis that maintaining continuity of opioid administration could be an effective therapeutic strategy to minimize the vulnerability to opioid use disorders.

## INTRODUCTION

Opioid use disorders are driven by both the rewarding euphoric states induced by opioids, and the subsequent negative dysphoric states induced by withdrawal and abstinence (Evans and Cahill, 2016; Koob, 2020). It is postulated that this leads to repeating cycles of negative reinforcement by alleviating withdrawal symptoms with continued opioid use (Evans and Cahill, 2016; Koob, 2020). Preclinical studies indicate that addiction-related behaviors are driven by both opioid-evoked and withdrawal-evoked adaptations in overlapping and distinct neural circuitry (Graziane et al., 2016; Hearing et al., 2016; Russell et al., 2016; Zhu et al., 2016; Madayag et al., 2019; McDevitt et al., 2019). Withdrawal is often studied in the context of short or long-term abstinence, but in the clinical setting, patients routinely experience repeated cycles of brief withdrawal periods – e.g., due to the pharmacokinetic variability of prescription opioid formulations purporting to provide continuous action (Ackerman et al., 2003). Therefore, it is also necessary to understand the adaptations in neural circuitry that occur in the context of repeated cycles of brief withdrawal from otherwise continuous opioid exposure.

In this study, we investigated how different patterns of opioid administration alter synaptic plasticity in the nucleus accumbens (NAc), utilizing our previously developed model of interrupting continuous morphine exposure with repeated bouts of naloxone-precipitated withdrawal (Lefevre et al., 2020). The premise for this model arises from previous studies demonstrating that intermittent patterns of opioid exposure cause sensitization of reward-related behaviors, whereas continuous opioid exposure induces tolerance (Shippenberg et al., 1988; Lett, 1989; Gaiardi et al., 1991; Shippenberg et al., 1996; Vanderschuren et al., 1997; Russo et al., 2007; Contet et al., 2008; Rothwell et al., 2010; Le Marec et al., 2011; Sun et al., 2014; Yu et al., 2014). To provide direct comparisons of patterns of opioid exposure, while controlling for confounding pharmacokinetic variables, we interrupted otherwise continuous morphine infusion with daily injections of naloxone (Lefevre et al., 2020). We found that the interruption of continuous morphine exposure with naloxone-precipitated withdrawal caused a reversal of psychomotor tolerance, augmented mesolimbic dopamine signaling, and induced striking transcriptional adaptations in the striatum (Lefevre et al., 2020). Thus this model provides a controlled method for studying how different patterns of opioid exposure affect reward circuitry (Cahill, 2020; Kibaly et al., 2021).

The NAc is a central hub in reward circuits and contains two subpopulations of medium spiny projection neurons (MSNs) that are classified by their expression of dopamine D1 receptors (D1-MSN) and dopamine D2 receptors (D2-MSN) (Le Moine and Bloch, 1995). Canonically, these two subpopulations are proposed to play opposing functional roles in the NAc, with D1-MSNs promoting reward and D2-MSNs negatively modulating reward (Hikida et al., 2010; Lobo and Nestler, 2011; Tai et al., 2012; Koo et al., 2014; Soares-Cunha et al., 2016; Cole et al., 2018; O’Neal et al., 2020; Soares-Cunha et al., 2020; O’Neal et al., 2022). Consistent with these canonical roles, NAc D1-MSNs were found to be activated in response to acute morphine, whereas NAc D2-MSNs are predominately activated in response to naloxone-precipitated morphine withdrawal (Enoksson et al., 2012).

Previous studies have shown that excitatory synaptic plasticity evoked by chronic morphine at D1-MSNs and D2-MSNs mediates rewarding and aversive behaviors, respectively (Graziane et al., 2016; Hearing et al., 2016; Russell et al., 2016; Zhu et al., 2016; Madayag et al., 2019; McDevitt et al., 2019). While there has been a large focus on excitatory synaptic plasticity, emerging evidence suggests chronic morphine administration alters inhibitory input onto MSNs, as well as the intrinsic excitability of MSNs (Koo et al., 2014; McDevitt et al., 2019). Though morphine exposure is known to modulate synaptic plasticity, it is not known how this is influenced by the pattern of opioid administration. Here, we sought to identify synaptic and cellular adaptations in NAc D1- and D2-MSNs following continuous versus interrupted morphine administration in male and female mice.

## MATERIALS AND METHODS

### Subjects

Experiments were performed with female and male mice maintained on a C57Bl/6J genetic background. Mice used in all experiments carried a single copy of a Drd1a-tdTomato BAC transgene mice (Shuen et al., 2008) and/or a Drd2-eGFP BAC transgene (Gong et al., 2003). Mice were 5-8 weeks old at the beginning of each experiment and housed in groups of 2-5 per cage, on a 12 hour light cycle (0600 – 1800h) at ~23°C with food and water provided ad libitum. Experimental procedures were approved by the Institutional Animal Care and Use Committee of the University of Minnesota.

### Drug Exposure

Morphine hydrochloride (Mallinckrodt) was dissolved in sterile saline (0.9%), and delivered continuously using osmotic minipumps (Alzet Model 2001) as previously described (Lefevre et al., 2020). Morphine concentration was adjusted for body weight to administer 63.2 mg/kg/day. Minipumps were filled with 300 μL of morphine or saline solution and primed overnight at 40°C. Once primed, minipumps were implanted under anesthesia (5% isoflurane/95% oxygen) through a small incision on the rump, which was then closed with wound clips. Carprofen (5 mg/kg, s.c.) was given as an analgesic before surgery and for 3 days following pump implantation or removal. Behavioral testing began 24 hours after pump implantation (Day 1), to allow for recovery from surgical anesthesia. To interrupt continuous morphine exposure, we injected mice with naloxone (Nlx; 10 mg/kg, s.c.) twice per day, with injections separated by a period of 2 h (Lichtblau and Sparber, 1981). Mice in the continuous morphine group received saline injections, and control groups implanted with saline pumps were injected with either saline or naloxone. In this study, the four experimental groups are thus described based on pump-injection treatment: the two saline pump groups are Sal-Sal or Sal-Nlx, and the two morphine pump groups are Mor-Sal (continuous) or Mor-Nlx (interrupted).

### Behavioral responses to morphine administration

We tested open-field locomotor activity in a clear plexiglass arena (ENV-510, Med Associates) housed within a sound-attenuating chamber, as previously described (Lefevre et al., 2020; Toddes et al., 2021). Mice were placed in the open field for 60 minutes on the first day after pump implantation (prior to the first set of saline or naloxone injections), in order to measure the acute effects of morphine infusion. This behavioral test was repeated in the same fashion prior to preparation of acute brain slices on day 6 (i.e., 24 hours after the last set of saline or naloxone injections).

### Electrophysiology

Parasagittal slices (240 μm) containing NAc shell were prepared using standard procedures (Pisansky et al., 2019; Toddes et al., 2021). This study focused on the NAc shell, as this subregion has been found to be more sensitive to opioid-evoked plasticity (Graziane et al., 2016; Hearing et al., 2016; Madayag et al., 2019; McDevitt et al., 2019). Mice were anesthetized with isoflurane and decapitated. Brains were quickly removed and placed in ice-cold cutting solution containing (in mM): 228 sucrose, 26 NaHCO_3_, 11 glucose, 2.5 KCl, 1 NaH_2_PO_4_-H_2_O, 7 MgSO_4_-7H_2_0, 0.5 CaCl_2_-2H_2_O. Slices were cut by adhering the lateral surface of the brain to the stage of a vibratome (Leica VT1000S), and then allowed to recover in a submerged holding chamber with artificial cerebrospinal fluid (aCSF) containing (in mM): 119 NaCl, 26.2 NaHCO_3_, 2.5 KCl, 1 NaH_2_PO_4_-H_2_O, 11 glucose, 1.3 MgSO_4_-7H_2_O, 2.5 CaCl_2_-2H_2_O. Slices recovered in warm ACSF (33°C) for 10-15 min and then equilibrated to room temperature for at least one hour before use. Slices were transferred to a submerged recording chamber and continuously perfused with aCSF at a rate of 2 mL/min at room temperature. All solutions were continuously oxygenated (95% O2/5% CO2).

Whole-cell recordings from MSNs were obtained under visual control using infrared-differential interference contrast optics on an Olympus BX51WI microscope. MSNs were identified by their morphology and hyperpolarized resting membrane potential (approximately −80 mV), with D1- and D2-MSNs differentiated by tdTomato or eGFP fluorophores, respectively. Recordings were performed using a MultiClamp 700B amplifier (Molecular Devices), filtered at 2 kHz, and digitized at 10 kHz. Data acquisition and analysis were performed online using Axograph software. Series resistance was monitored continuously, and experiments were discarded if resistance changed by >20%.

For current clamp recordings, borosilicate glass electrodes (3-5 MΩ) were filled with (in mM): 120 K-Gluconate, 20 KCl, 10 HEPES, 0.2 EGTA, 2 MgCl_2_, 4 ATP-Mg, 0.3 GTP-Na (pH 7.2-7.3). Before commencing recordings, MSN resting membrane potentials were adjusted to approximately −80mV. Cells were injected with a series of current steps (800 ms duration) from −160 to +260 pA, with a 20pA step increment. Maximum firing rate was calculated as the number of spikes over the 800 ms step that could be sustained without inducing a depolarization block, and averaged from 3 cycles of current steps. Rheobase was calculated as the minimum current injection required to induce action potential firing. The current-voltage (IV) relationship, used to measure ‘steady-state’ membrane properties, was obtained by measuring voltage response at subthreshold current steps (−160 to +60pA) (Kourrich and Thomas, 2009; Pisansky et al., 2019).

Voltage-clamp recordings were made with borosilicate glass electrodes (2–5 MΩ) filled with the following (in mM): 120 CsMeSO_4_, 15 CsCl, 10 TEA-Cl, 8 NaCl, 10 HEPES, 1 EGTA, 5 QX-314, 4 ATP-Mg, and 0.3 GTP-Na, pH 7.2–7.3. Spermine (0.1 mM) was included when measuring the current-voltage relationship of AMPAR currents. All reported holding currents are corrected for a liquid junction potential of ~10 mV. Excitatory and inhibitory evoked currents were electrically stimulated using a glass monopolar electrode filled with aCSF (ISO-flex, AMPI) at 0.1Hz. Spontaneous excitatory post-synaptic currents (sEPSCs) were recorded at a holding potential of −80 mV and pharmacologically isolated using GABA_A_ receptor antagonist picrotoxin (50 μM).Spontaneous inhibitory post-synaptic currents (sIPSCs) were recorded at a holding potential of 0 mV and pharmacologically isolated using NMDA and AMPA receptor antagonists, D-APV (50 μM) and NBQX (10 μM), respectively. For all spontaneous recordings, at least 200 events per cell were acquired in 15 s blocks and detected using a threshold of 5 pA; all events included in the final data analysis were verified by eye.

For AMPAR/NMDAR ratio experiments, recordings were made in the presence of picrotoxin and MSNs were voltage clamped at +40 mV, whereby NMDARs are no longer blocked by magnesium. Once a stable dual-component excitatory baseline was acquired, D-APV (50 μM) was bath applied to isolate the AMPA receptor-mediated current. The NMDA receptor mediated current was obtained by digital subtraction of the AMPAR current from the dual-component current, and peak amplitudes were used to calculate the ratio. Paired-pulse ratios (PPR) were acquired in the presence of picrotoxin, by voltage-clamping MSNs at −80 mV and electrically evoking two EPSCs of equal intensity with a 25 ms inter-stimulus interval, and then calculating the ratio between the peak amplitude of the second EPSC and the amplitude of the first EPSC. To investigate AMPAR subunit composition, we conducted an I-V analysis of pharmacologically isolated AMPAR EPSCs at holding potentials of −80, −40, 0, +20 and +40 mV, and normalized peak amplitudes to −80 mV. The peak EPSC amplitude at +40 mV to −80 mV was divided to calculate the rectification index. For excitatory/inhibitory (E/I) ratio experiments, MSNs were voltage clamped at −80 mV, and both excitatory and inhibitory presynaptic afferents were electrically stimulated without pharmacological isolation. Once a stable dual-component current was acquired, picrotoxin (50 μM) was bath applied to isolate the excitatory AMPA receptor-mediated current. The inhibitory GABA receptor mediated current obtained by digital subtraction of the excitatory current from the dual-component current, and peak amplitudes were used to calculate the ratio. Decay time constants for the NMDAR EPSC (+40 mV), AMPAR EPSC (−80 mV), and GABAR IPSC (−80 mV) were derived from averaged currents by fitting to double exponential equations using Easy Electrophysiology software, and the weighted mean decay time constants were calculated as previously described (Rumbaugh and Vicini, 1999).

### Experimental design and statistical analyses

All experiments were conducted using hemizygous BAC transgenic mice on a C57Bl/6J background (Nelson et al., 2012), to avoid potentially detrimental effects of breeding a randomly inserted BAC transgene to homozygosity (Kramer et al., 2011). Similar numbers of male and female mice were used in all experiments, with sample size indicated in figure legends. Sex was included as a variable in factorial ANOVA models analyzed using IBM SPSS Statistics v24, with repeated measures on within-subject factors. Kolmogorov-Smirnov tests were conducted using GraphPad Prism 7. All summary data are displayed as mean + SEM, with individual data points from male and female mice shown as closed and open symbols, respectively. Significant interactions were decomposed by analyzing simple effects (i.e., the effect of one variable at each level of the other variable). Significant main effects were analyzed using Tukey’s post-hoc tests. The Type I error rate was set to α = 0.05 (two-tailed) for all comparisons.

## RESULTS

### Divergent behavioral adaptations induced by continuous versus interrupted morphine exposure

We previously demonstrated that psychomotor tolerance develops during continuous morphine administration, but can be reversed by daily interruption of opioid exposure with naloxone-precipitated withdrawal (Lefevre et al., 2020). In this study, we used the same model to characterize adaptations in intrinsic excitability and synaptic plasticity in NAc MSNs. All mice received subcutaneous implantation of an osmotic minipump that delivered continuous infusion of morphine (63.2 mg/kg/day) or saline over the course of six days (Figure 1A). Using a factorial design, we also administered twice-daily injections of saline or naloxone (10 mg/kg) on days 1-5 after pump implantation, so that opioid administration was either “continuous” (morphine pump plus saline injections) or “interrupted” (morphine pump plus naloxone injections). As previously described (Lefevre et al., 2020), the Mor-Nlx group showed weight loss following each day of naloxone-precipitated withdrawal (Table 1), and the magnitude of this effect showed a significant linear increase over days (F_1,35_ = 13.34, p = 0.001).

**Figure 1.**
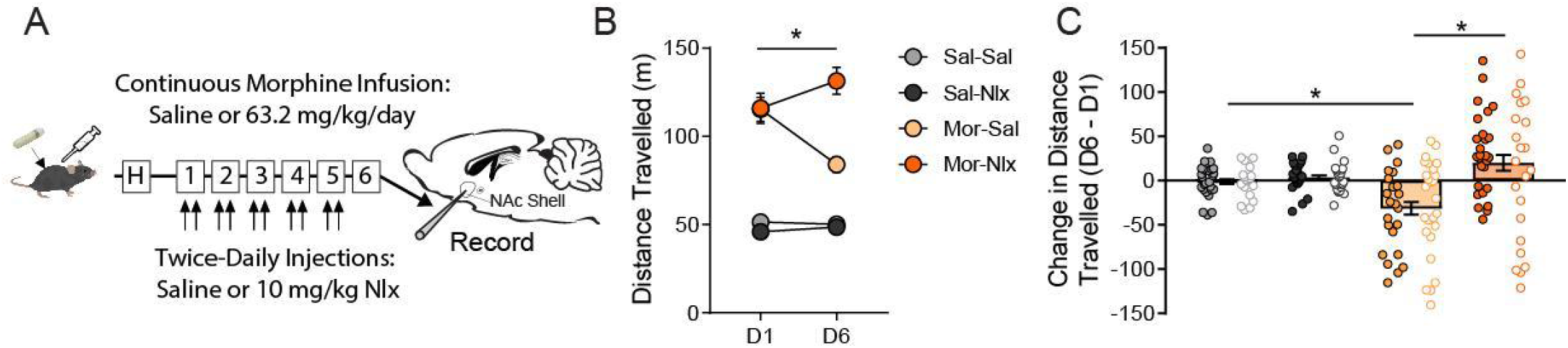
Interruption of continuous morphine exposure with daily naloxone injections. ***A*,** Experimental timeline including one habituation day (“H”), followed by pump implantation providing continuous infusion of morphine or saline for 5 days, interrupted by twice-daily injections (separated by 2 h) of saline or naloxone. Locomotor activity was recorded on Day 1 prior to injections and on Day 6. *Ex-vivo* sagittal slices containing the NAc shell for electrophysiology recordings were collected immediately following behavioral testing on Day 6. ***B*,** Locomotor activity on the first (D1) and last (D6) day of exposure (n = 45-51/group). ***C*,** Change in locomotor activity on D6 versus D1, depicted for individual mice at each dose. All groups contained similar numbers of female mice (open symbols) and male mice (closed symbols). *p < 0.05 between groups, Tukey post-hoc test.

**Table 1.** Percentage of body weight lost following daily injections of saline or naloxone.

|  | Sal-Sal (36) | Sal-Nlx (38) | Mor-Sal (40) | Mor-Nlx (38) |
| --- | --- | --- | --- | --- |
| Day 1 | 0.32 ± 0.26% | -0.66 ± 0.24% | 0.15 ± 0.24% | -4.15 ± 0.33% |
| Day 2 | -0.45 ± 0.19% | -0.58 ± 0.15% | -0.76 ± 0.33% | -5.16 ± 0.30% |
| Day 3 | -0.63 ± 0.19% | -0.65 ± 0.23% | 0.13 ± 0.21% | -5.86 ± 0.31% |
| Day 4 | -0.48 ± 0.17% | -0.69 ± 0.18% | -0.32 ± 0.25% | -5.93 ± 0.47% |
| Day 5 | 0.08 ± 0.24% | -0.40 ± 0.23% | -0.42 ± 0.18% | -5.99 ± 0.37% |

*All data are presented as mean ± SEM; number in parenthesis represents sample size for each treatment group*

Mice were placed in an open field activity chamber on days 1 and 6 after pump implantation, to measure distance traveled (Figure 1B). On day 1, prior to the first injection of saline or naloxone, mice with morphine pumps displayed a robust increase in psychomotor activation. On day 6, psychomotor tolerance was observed in the continuous morphine group, whereas this behavioral effect was reversed after interrupted morphine treatment (Morphine × Naloxone × Day interaction: F_1,189_ = 10.38, p = 0.001). There was also a main effect of Sex (F_1,185_ = 4.0, p = 0.047), indicating that overall locomotor activity was higher in females compared to males. However, there were no significant interactions between Sex and any other factor. Changes in locomotor activity over time were most apparent after computing the change in distance traveled between day 1 and day 6 for each individual animal (Figure 1C). These patterns of behavioral adaptation produced by continuous and interrupted morphine exposure are consistent with our previous report (Lefevre et al., 2020).

### Intrinsic excitability is not altered by either continuous or interrupted morphine exposure

MSNs in the NAc express opioid receptors (Charbogne et al., 2017), and MSN intrinsic excitability can be modulated by acute activation of opioid receptors (Ma et al., 2012; Trieu et al., 2022). It has also been shown that repeated morphine exposure alters the intrinsic excitability of NAc MSNs (Heng et al., 2008; McDevitt et al., 2019). Thus, we sought to characterize intrinsic firing properties of NAc MSNs following continuous or interrupted morphine administration. We used whole-cell current clamp recordings to measure the number of spikes elicited by a series of depolarizing current injections. Neither pattern of morphine exposure altered the intrinsic excitability of either D1-MSNs (Figure 2A-B) or D2-MSNs (Figure 2C-D). There were also no changes in passive membrane properties or rheobase (Figure 2E-F). We also analyzed qualitative characteristics of the spike trains, including latency to first spike, train duration and inter-spike interval (Figures 2G-L). There were no changes in these spike train characteristics in any treatment group. These data suggest that neither continuous nor interrupted morphine exposure alter the intrinsic excitability of NAc MSNs at the time point examined.

**Figure 2.**
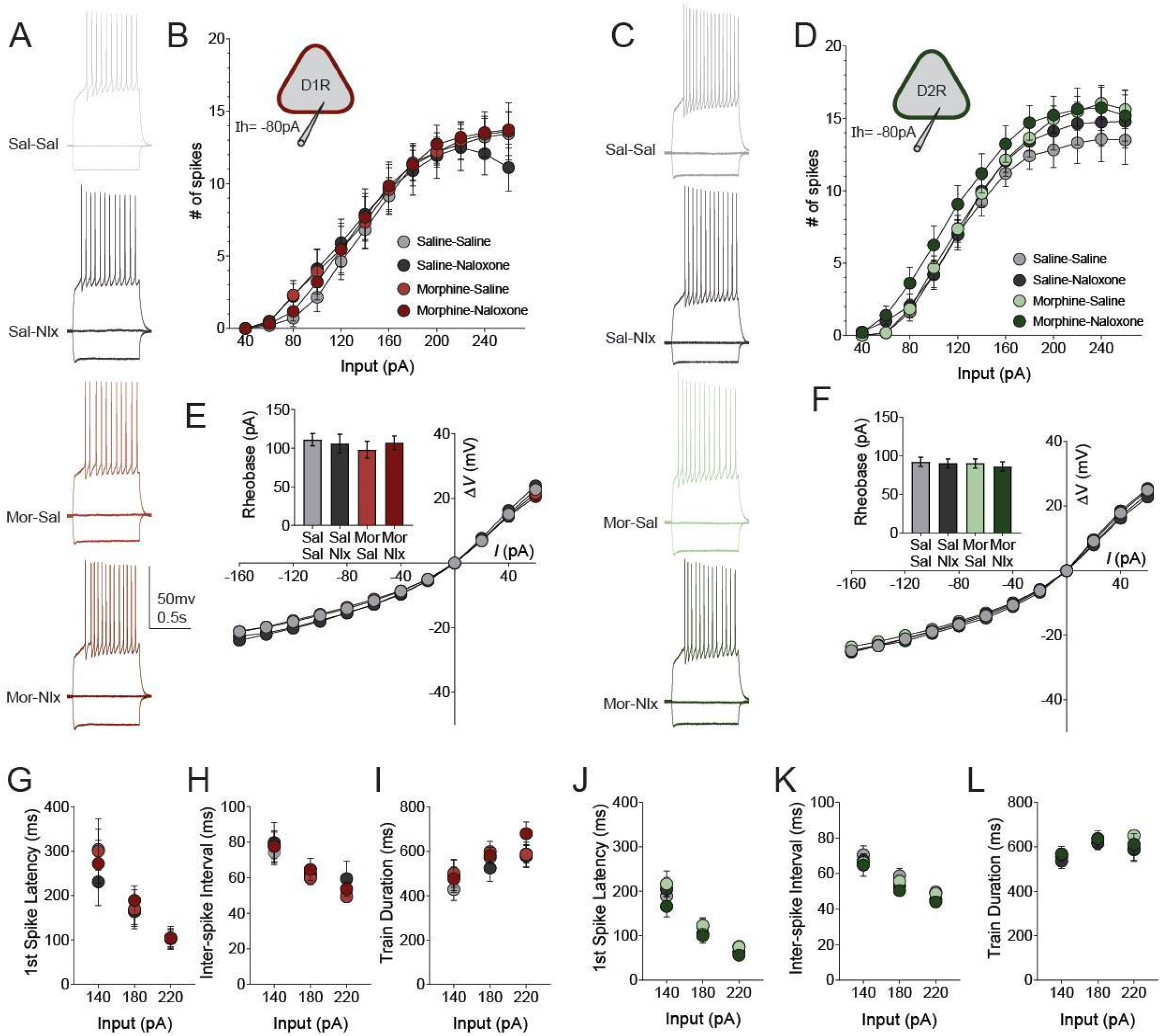
Whole-cell current-clamp electrophysiology recordings from MSNs in the NAc shell to measure intrinsic excitability. ***A, C*,** Representative traces at currents steps −160pA, 0pA and +180pA in D1-MSNs (*n* = 9-13/group) (***A***) and D2-MSNs (*n* = 17-23/group) (***C***). ***B, D*,** Positive current steps incrementing in 20pA from +40pA to +260pA was used to calculate the number of action potential ‘spikes’ that could be induced over the 800 ms pulse in D1- (***B***) and D2-MSNs (***D***). ***E, F*,** Subthreshold current steps (800 ms) were used to plot the current-voltage (IV) relationship and found no morphine-induced changes in passive membrane properties. Rheobase was calculated as the threshold current injection required to induce action potential firing (inset). ***G-L*,** MSN firing properties at representative levels of depolarizing current injection (140, 180 and 220pA). Latency to the first action potential in D1- (***G***) and D2-MSNs (***J***). Inter-spike interval in D1- (***H***) and D2-MSNs (***K***). Train duration D1- (***I***) and D2-MSNs (***L***). No main effects of sex or drug treatment were identified.

### Continuous and interrupted morphine exposure cause divergent changes in spontaneous excitatory currents

A number of previous studies have shown that daily morphine injections (which cause psychomotor sensitization) produce cell type-specific changes in excitatory synaptic input to D1-MSNs and D2-MSNs (Graziane et al., 2016; Hearing et al., 2016; Zhu et al., 2016; Madayag et al., 2019; McDevitt et al., 2019). We sought to identify whether continuous morphine exposure (which causes psychomotor tolerance) would produce distinct adaptations in excitatory transmission, and if different adaptations would develop after interruption of continuous morphine by daily naloxone injections. We first measured spontaneous excitatory post-synaptic currents (sEPSCs) from D1- and D2-MSNs (Figure 3A-B). In the NAc shell, sEPSC amplitude was significantly higher in the D1-MSNs of Mor-Nlx mice compared to Mor-Sal mice (Figure 3C-D; main effect of Group: F_3,57_ = 3.44, p = 0.023). A similar trend was observed in D2-MSNs (Figure 3E-F; main effect of Group: F_3,58_ = 2.49, p = 0.069). No significant main effects on sEPSC frequency were observed in D1-MSNs. However, Mor-Nlx mice showed a higher distribution of longer inter-event intervals (IEIs) in D1-MSNs relative to Sal-Nlx controls (Figure 3H; KS test, D = 0.18, p = 0.003). This change in the frequency of events may reflect the increase in Sal-Nlx group, as opposed to a decrease in frequency events evoked by Mor-Nlx. Interestingly, there was a main effect of Sex on sEPSC frequency in D2-MSNs, with males overall showing a higher frequency of events than females (F_1,58_ = 5.50, p = 0.022).

**Figure 3.**
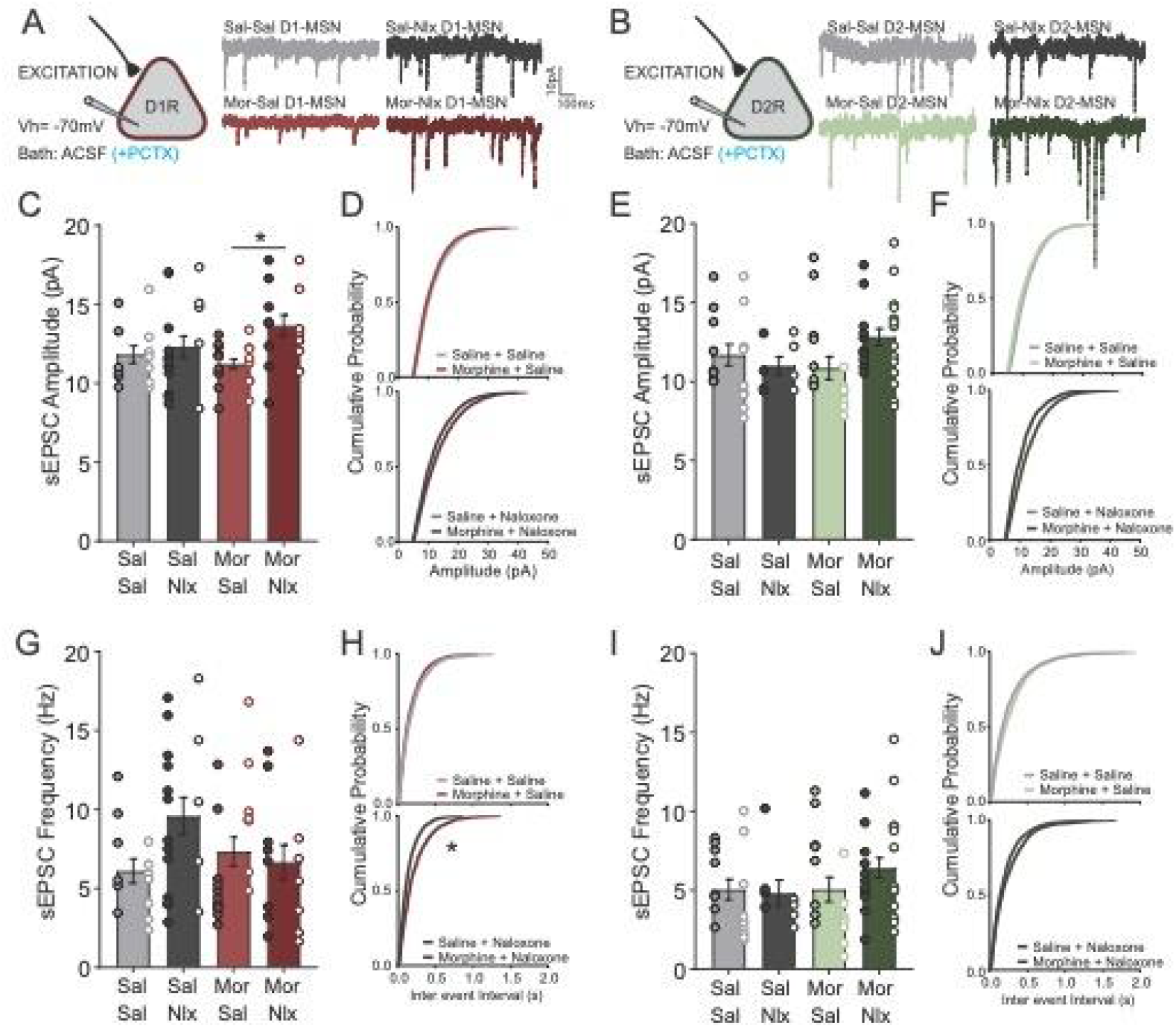
Electrophysiological recordings from MSNs in the NAc to assess spontaneous excitatory synaptic transmission. ***A-B*,** Schematic diagram showing whole-cell voltage clamp recordings held at −70 mV from MSNs identified by the expression of Drd1-tdTomato (***A***) or Drd2-eGFP (***B***). Example traces show sEPSCs recorded for Sal-Sal (D1, n=13 cells; D2, n=17 cells), Sal-Nlx (D1, n=18 cells; D2, n=8 cells), Mor-Sal (D1, n=19 cells; D2, n=16 cells), Mor-Nlx (D1, n=15 cells; D2, n=25 cells). Scale bar applies to all traces. ***C-F*,** Average sEPSC amplitude and cumulative probability plots for D1-MSNs (***C, D***) and D2-MSNs (***E, F***) separated by injection (Sal vs. Nlx). ***G-J*,** Average sEPSC frequency and cumulative probability plots for D1-MSNs (***G, H***) and D2-MSNs (***I, J***) separated by injection (Sal vs. Nlx). Male mice are represented by closed symbols and female mice are represented by open symbols. \**p*<0.05 according to LSD *post-hoc* test (***C***) or Kolmogorov-Smirnov test (***H***).

### Convergent effects of continuous and interrupted morphine exposure on evoked excitatory transmission

We next investigated whether continuous and interrupted morphine exposure altered excitatory synaptic strength, by measuring the ratio of AMPA receptor mediated current to NMDA receptor mediated current (i.e. the AMPAR/NMDAR ratio; Figure 4A). In D1-MSNs, the AMPAR/NMDAR ratio was significantly decreased in both morphine groups in males, but not females (Figure 4B; Pump x Sex interaction: F_1,33_ = 4.51, p = 0.041). No significant changes in the AMPAR/NMDAR ratio were found in D2-MSNs (Figure 4C). We also did not detect changes in the coefficient of variation for evoked AMPAR EPSCs (Table 2) or the decay kinetics of NMDAR ESPCs (Table 2) in any treatment group or cell type.

**Figure 4.**
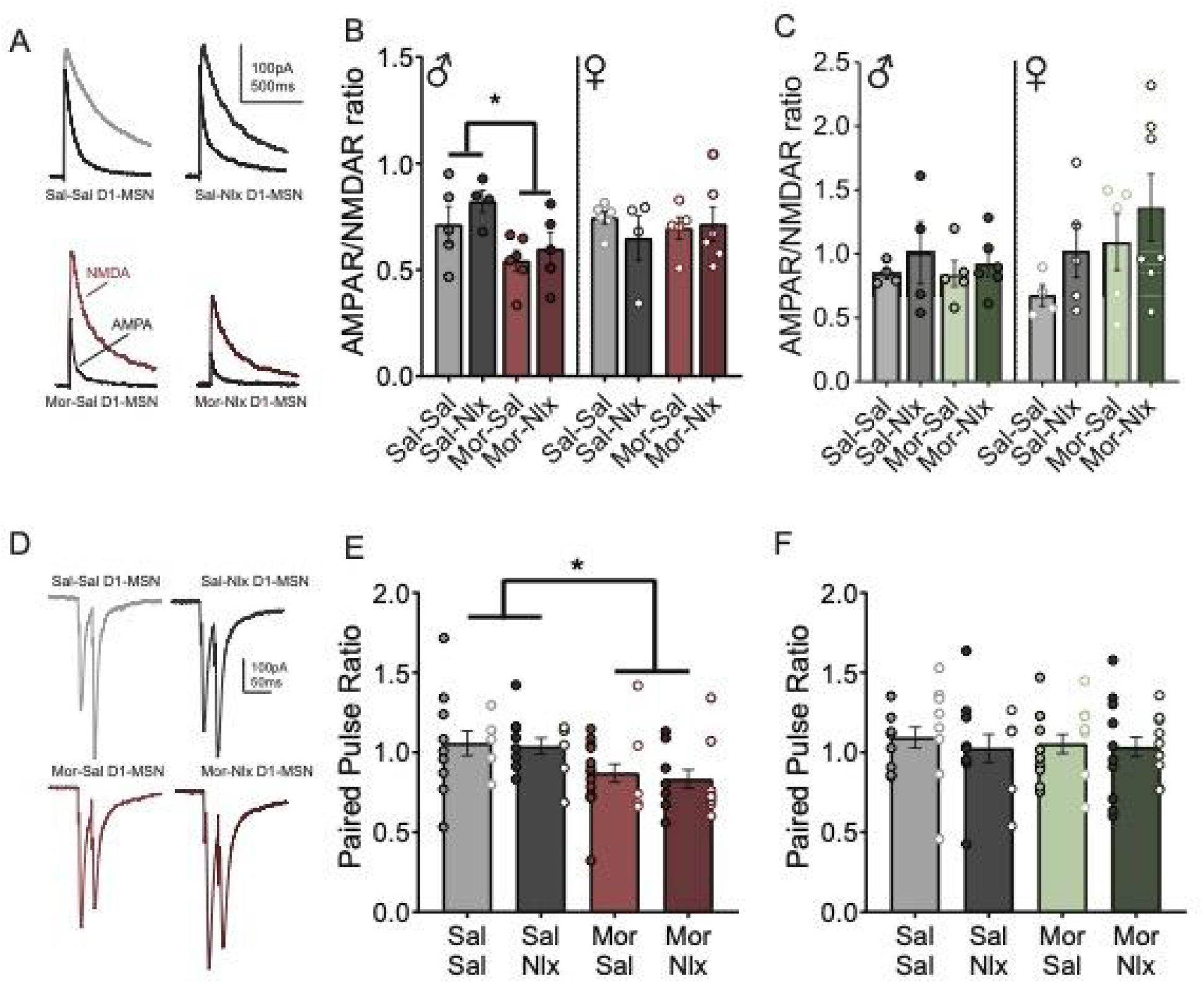
Morphine induced changes in NAc MSN evoked excitatory synaptic transmission. ***A-C*,** Excitatory synaptic strength was assessed as the evoked AMPAR/NMDAR ratio in whole-cell voltage clamped (+40 mV) MSNs. ***A*,** Example AMPAR and NMDAR traces from D1-MSNs in male mice. Averaged AMPAR/NMDAR ratios are shown from D1- (***B***) and D2-MSNs (***C***) separated by sex (*n* = 4-7/sex/group). ***D-F*,** Probability of presynaptic glutamate release was measured by the ratio of paired-pulse evoked EPSCs (25ms apart) in whole-cell voltage clamped (−80 mV) MSNs. ***D*,** Representative traces of paired pulse ratio from D1-MSNs. Mean paired pulse ratio in D1- (***E***) and D2-MSNs (***F***), pooled for sex (*n* =13-18/group). \**p*<0.05 Main effect of Morphine.

**Table 2:** Properties of evoked AMPAR and NMDAR EPSCs measured at +40 mV.

|  |  | Sal-Sal | Sal-Nlx | Mor-Sal | Mor-Nlx |
| --- | --- | --- | --- | --- | --- |
| D1-MSN | AMPA RI | 0.44 ± 0.08 (5) | 0.62 ± 0.04 (5) | 0.52 ± 0.02 (5) | 0.46 ± 0.05 (8) |
|  | AMPA CV | 0.12 ± 0.02 (11) | 0.12 ± 0.02 (8) | 0.12 ± 0.02 (11) | 0.12 ± 0.01 (11) |
|  | NMDAR decay | 207.38 ± 30.10 (10) | 214.71 ± 23.28 (7) | 231.86 ± 23.08 (7) | 214.01 ± 9.35 (9) |
| D2-MSN | AMPA RI | 0.67 ± 0.07 (7) | 0.50 ± 0.07 (6) | 0.59 ± 0.04 (6) | 0.58 ± 0.03 (4) |
|  | AMPA CV | 0.12 ± 0.02 (8) | 0.10 ± 0.01 (9) | 0.13 ± 0.01 (10) | 0.14 ± 0.02 (12) |
|  | NMDAR decay | 248.08 ± 27.57 (7) | 289.32 ± 60.00 (7) | 277.47 ± 40.51 (9) | 269.76 ± 49.39 (8) |

All data are expressed as mean ± SEM; number in parenthesis represents sample size. Abbreviations: RI = rectification index, CV = coefficient of variance.

We used the paired-pulse ratio (PPR) to assess changes in presynaptic glutamate release (Figure 4D). In D1-MSNs, PPR was significantly decreased in both morphine groups, to a comparable extent in both sexes (Figure 4E; main effect of Pump F_1,52_ = 7.59, p = 0.008). However, PPR was not affected in D2-MSNs (Figure 4F). These would suggest an increase in glutamate release probability onto D1-MSNs, as previously reported following other patterns of morphine exposure (Hearing et al., 2016). Calcium-permeable AMPARs (CP-AMPARs) are often recruited during periods of drug-evoked synaptic plasticity (Luscher and Malenka, 2011; Wolf and Tseng, 2012; Yuan and Bellone, 2013), and their synaptic incorporation leads to inward rectification of the current-voltage (IV) relationship for evoked AMPAR currents. However, we did not detect changes in the rectification index in any treatment group or cell type (Table 2).

### Divergent changes in spontaneous inhibitory transmission after continuous and interrupted morphine exposure

The functional output of NAc MSNs is tightly regulated by synaptic input from a small population (~5%) of local inhibitory interneurons (Sesack and Grace, 2010), as well as lateral inhibitory connections between MSNs (Burke et al., 2017). Previous reports indicate that chronic morphine administration also alters inhibitory transmission onto NAc MSNs (Koo et al., 2014; McDevitt et al., 2019), and we have previously shown that interrupted and continuous morphine administration cause differential changes in the expression of GABA receptor subunits (Lefevre et al., 2020). To assess if these divergent transcriptional adaptations conferred functional alterations in inhibitory synaptic transmission, we measured spontaneous inhibitory post-synaptic currents (sIPSC). In D1-MSNs, we noted a trend toward a main effect of Group (Figure 5A; F_3,68_ = 2.71, p = 0.052) on sIPSC amplitude, with Mor-Sal mice showing an increase compared to Sal-Sal controls (p < 0.05). For sIPSC frequency, there was a main effect of Sex in D1-MSNs (F_1,68_ = 4.41, p = 0.039) with males showing a higher frequency than females. Together with the sex differences reported in Figure 3C, this indicates that males have higher inhibitory input onto D1-MSNs and a higher excitatory input onto D2-MSNs than their female counterparts. Two-way ANOVA revealed no significant effects on frequency in D2-MSNs, however the Kolmogorov-Smirnov test (Figure 5D, inset) identified a significant shift in the cumulative distribution of inter-event-intervals (IEIs). Mor-Nlx mice showed a higher distribution of longer IEIs relative to Sal-Nlx controls, indicative of a decrease in the frequency of events. These results suggest that continuous morphine administration increases inhibitory signaling selectively onto D1-MSNs, likely via a post-synaptic mechanism. In contrast, interrupted morphine may decrease inhibitory input selectively onto D2-MSNs.

**Figure 5.**
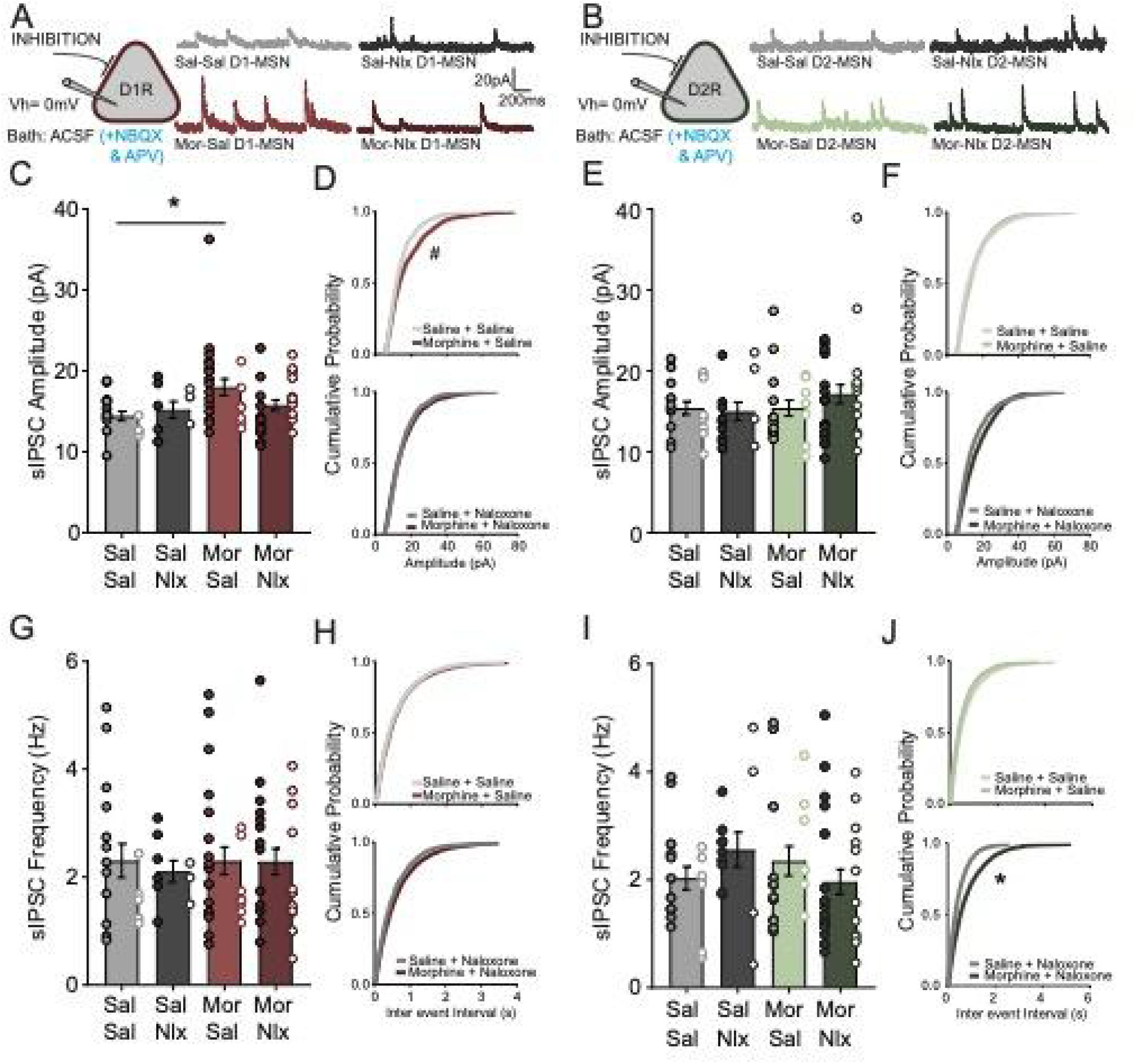
Electrophysiological recordings from MSNs in the nucleus accumbens to assess inhibitory synaptic transmission. ***A, B*,** Schematic diagram showing whole-cell voltage clamp recordings held at 0mV from MSNs identified by the expression of Drd1-tdTomato (***A***) or Drd2-eGFP (***B***). Example traces show sIPSC recorded for Sal-Sal (D1, n=17 cells; D2, n=18 cells), Sal-Nlx (D1, n=9 cells; D2, n=13 cells), Mor-Sal (D1, n=24 cells; D2, n=20 cells), Mor-Nlx (D1, n=26 cells; D2, n=29 cells). ***C-F*,** Average sIPSC amplitude and cumulative probability plots for D1-MSNs (***C, D***) and D2-MSNs (***E, F***) separated by injection (Sal vs. Nlx). ***G-J*,** Average sIPSC frequency and cumulative probability plots for D1-MSNs (***G, H***) and D2-MSNs (***I, J***) separated by injection (Sal vs. Nlx). Male mice are represented by closed symbols and female mice are represented by open symbols. #*p*=0.052 according to Kolmogorov-Smirnov test (***D***). \**p*<0.05 according to LSD *post-hoc* test (***C***) or Kolmogorov-Smirnov test (***J***).

### Divergent morphine induced adaptations in excitatory to inhibitory transmission

Since we identified morphine-induced changes in both sIPSCs and sEPSCs, we sought to identify whether this would be reflected as a shift in the balance between excitatory and inhibitory synaptic input. To assess this possibility, we directly measured the ratio between currents mediated by AMPA receptors and GABA receptors in MSNs voltage-clamped at −80 mV (Figure 6A-B). The ratio of peak AMPA receptor to peak GABA receptor-mediated currents (i.e. excitation/inhibition ratio) in D1-MSNs was significantly reduced in male Mor-Sal mice compared their female Mor-Sal counterparts (Figure 6C; Two-way ANOVA; Group x Sex F_3,30_ = 3.89, p = 0.018). Together with the decrease in AMPAR/NMDAR ratio observed in Mor-Sal males (Figure 4A), this would suggest that AMPAR-mediated transmission is decreased specifically in male Mor-Sal treated mice. No significant changes in AMPAR/GABAR ratio were identified in D2-MSNs (Figure 6D).

**Figure 6.**
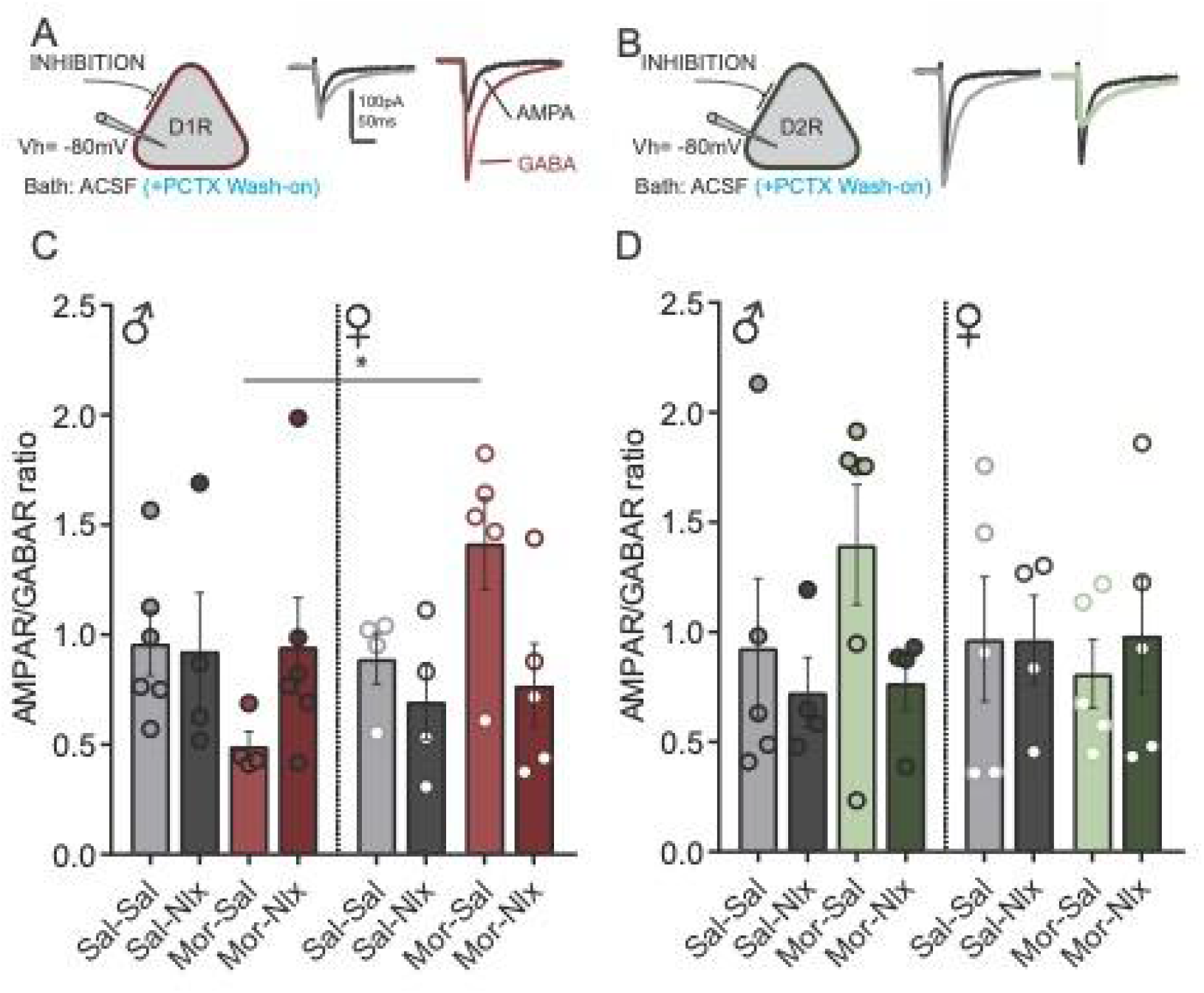
Evoked ratio of excitatory to inhibitory transmission in NAc MSNs. ***A-B*,** Schematic diagram showing whole-cell voltage clamp recordings from MSNs held at −80mV. Representative traces from male Sal-Sal and Mor-Sal treated mice showing AMPAR- and GABAR-mediated currents on D1-MSNs (***A***) and D2-MSNs (***B***). ***C-D*,** Average AMPAR/GABAR ratio plots for D1-MSNs (***C***) and D2-MSNs (***D***) separated by sex (*n* = 4-6/sex/group). \**p*<0.05 according to LSD *post-hoc* test.

We also analyzed the decay kinetics of both the AMPAR EPSCs and GABAR IPSCs, and consistent with the selective changes observed in AMPAR/GABAR ratio, found significant adaptations in D1- but not D2-MSNs (Table 3). For the decay kinetics of AMPAR EPSCs in D1-MSNs, we identified a significant Group x Sex interaction (F_3,26_ = 4.32, p = 0.013). In female D1-MSNs, there was a significant difference between Mor-Sal and Mor-Nlx mice, whereas in the male mice there was a significant difference between Sal-Nlx and Mor-Nlx groups. For the decay kinetics of GABAR IPSCs in D1-MSNs, we also identified a significant Group x Sex interaction (F_3,26_ = 3.28, p = 0.037). This interaction was driven by differences between the Sal-Sal and Mor-Sal treatment group (Morphine x Sex interaction: F_1,17_ = 9.17, p = 0.010). Males in the Mor-Sal group showed significantly faster decay of GABAR IPSCs compared to Sal-Sal males (F_1,8_ = 5.92, p = 0.041), whereas females in the Mor-Sal group tended to have slower decay compared to Sal-Sal females (F_1,5_ = 3.48, p = 0.12), correlating with the changes observed in the AMPAR/GABAR ratio (Figure 6C). These concomitant changes in male D1-MSNs could be due to changes in GABAR subunit composition that increase peak amplitude and rate of decay, or a redistribution of inhibitory synaptic inputs that favors more proximal sites near the cell body. Either of these changes could also explain the increased sIPSC amplitude caused by Mor-Sal treatment in male D1-MSNs (Figure 5C), but surprisingly, female D1-MSNs appear to show a similar increase in sIPSC amplitude caused by Mor-Sal treatment. Together, these results point to a unique forms and mechanisms of D1-MSN inhibitory synaptic plasticity caused by continuous morphine treatment in each sex, while also indicating these forms of plasticity are not present when continuous morphine exposure is repeatedly interrupted by naloxone-precipitated withdrawal.

**Table 3:** Properties of evoked AMPAR EPSCs and GABAR IPSCs measured at −80 mV.

|  |  | Sex | Sal-Sal | Sal-Nlx | Mor-Sal | Mor-Nlx |
| --- | --- | --- | --- | --- | --- | --- |
| D<br>1<br>-<br>M<br>S<br>N | AMPA<br>decay | *Male* | 23.16 ± 2.22 (6) | 17.23 ± 1.57 (3)* | 24.54 ± 1.64 (4) | 28.14 ± 2.37 (5)* |
|  |  | *Female* | 17.17 ± 2.82 (4) | 18.02 ± 2.90 (4) | 26.15 ± 1.19 (3)* | 15.66 ± 1.39 (5)* |
|  | GABAR<br>decay | *Male* | 65.50 ± 5.84 (6)* | 45.06 ± 5.20 (3) | 45.58 ± 4.58 (4)* | 52.26 ± 4.75 (5) |
|  |  | *Female* | 46.81 ± 9.78 (4) | 52.86 ± 9.81 (4) | 79.62 ± 15.8 (3) | 48.88 ± 10.0 (5) |
| D<br>2<br>-<br>M<br>S<br>N | AMPA<br>decay | *Male* | 20.24 ± 2.02 (5) | 19.87 ± 1.76 (4) | 26.32 ± 3.28 (6) | 19.80 ± 3.26 (4) |
|  |  | *Female* | 23.89 ± 3.30 (5) | 23.36 ± 2.59 (4) | 27.90 ± 4.61 (5) | 23.65 ± 3.26 (5) |
|  | GABAR<br>decay | *Male* | 49.26 ± 6.44 (5) | 54.14 ± 8.47 (4) | 71.66 ± 16.2 (6) | 58.14 ± 11.15 (4) |
|  |  | *Female* | 66.77 ± 12.5 (5) | 57.56 ± 7.21 (4) | 64.14 ± 14.3 (5) | 64.60 ± 9.09 (5) |

All data are expressed as mean ± SEM; number in parenthesis represents sample size. \**p* < 0.05 between treatment groups within sex.

## DISCUSSION

In this this study, we identified cell type-specific and sex-dependent changes in excitatory and inhibitory synaptic plasticity within the NAc shell induced by continuous versus interrupted morphine administration (Table 4). Consistent with our previous behavioral findings (Lefevre et al., 2020), we showed that interruption of otherwise continuous morphine administration with daily naloxone injections reversed the psychomotor tolerance associated with continuous morphine exposure (Figure 1). However, we did not detect changes in the intrinsic excitability of NAc MSNs following either pattern of morphine administration. We observed some adaptations in excitatory synaptic function that were evoked by both patterns of morphine administration, but other adaptations at excitatory and inhibitory synapses were unique to only one pattern of morphine administration. Our data provide further evidence that the pattern of opioid administration dictates drug-evoked adaptations in the mesolimbic system.

**Table 4:** Summary of electrophysiology measures following continuous or interrupted morphine.

| Experimental Measure |  | Continuous Morphine |  |  |  | Interrupted Morphine |  |  |  |
| --- | --- | --- | --- | --- | --- | --- | --- | --- | --- |
|  |  | D1-MSN |  | D2-MSN |  | D1-MSN |  | D2-MSN |  |
|  |  | ♂ | ♀ | ♂ | ♀ | ♂ | ♀ | ♂ | ♀ |
| sEPSC | Amplitude | ↓ | ↓ |  |  | ↑ | ↑ |  |  |
|  | Frequency |  |  |  |  |  |  |  |  |
| A/N ratio | Synaptic Strength | ↓ |  |  |  | ↓ |  |  |  |
| PPR | Glutamate release probability | ↓ | ↓ |  |  | ↓ | ↓ |  |  |
| sIPSC | Amplitude | ↑ | ↑ |  |  |  |  |  |  |
|  | Frequency |  |  |  |  |  |  | ↓ | ↓ |
| A/G ratio | Excitatory/Inhibitory Balance | ↓ | ↑ |  |  |  |  |  |  |
| Spike # | Intrinsic Excitability |  |  |  |  |  |  |  |  |

### Continuous and interrupted morphine do not alter MSN intrinsic excitability

The functional output of NAc MSNs is determined by the integration of their intrinsic membrane and synaptic properties (O’Donnell et al., 1999). Alterations in NAc MSN intrinsic membrane excitability can be caused by exposure to psychostimulants (Zhang et al., 1998; Dong et al., 2006; Kourrich and Thomas, 2009; Mu et al., 2010; Wang et al., 2018), as well as opioids (Heng et al., 2008; McDevitt et al., 2019). Furthermore, NAc MSNs express the mu opioid receptor, and their intrinsic excitability can be directly modulated by mu opioid receptor activation (Ma et al., 2012; Trieu et al., 2022). However, intrinsic membrane excitability of NAc MSNs was not altered on day 6 following either continuous or interrupted morphine administration (Figure 2). A number of technical differences including species, pattern of opioid administration/withdrawal, anatomical region, and cell type-specificity may explain the discrepancy with a previous report that repeated morphine administration decreased intrinsic excitability of MSNs (Heng et al., 2008). However, in a technically similar study, McDevitt et al. (2019) identified increased intrinsic excitability specifically in D2-MSNs of the NAc shell in male and female mice after only 1 day of withdrawal from repeated morphine injections (McDevitt et al., 2019). This suggests that sustained withdrawal from morphine that lasts at least 24 hours may be necessary to induce adaptations in NAc MSN intrinsic excitability.

### Convergent excitatory synaptic plasticity following continuous and interrupted morphine administration

Converging excitatory inputs to NAc shell MSNs play a critical role in both opioid reward and the aversive effects of opioid withdrawal (Hearing et al., 2016; Russell et al., 2016; Zhu et al., 2016). Previous studies report that at early withdrawal, chronic morphine-evoked increased excitatory input to D2-MSNs was critical for the expression of aversive behaviors, with no changes observed at excitatory inputs to D1-MSNs (Graziane et al., 2016; Russell et al., 2016; Zhu et al., 2016; Madayag et al., 2019; McDevitt et al., 2019). Following extended abstinence from chronic morphine administration, excitatory inputs to D1-MSNs are strengthened and mediate reward-related behaviors (Hearing et al., 2016; Madayag et al., 2019), whereas excitatory input to D2-MSNs is weakened (Graziane et al., 2016; Hearing et al., 2016). In the current study, we prepared acute brain slices prior to spontaneous withdrawal from continuously delivered morphine, to directly assess the effects of repeated daily cycles of naloxone-precipitated withdrawal on excitatory synaptic plasticity. Perhaps surprisingly, at this time point we identified excitatory synaptic adaptations specifically onto D1-MSNs but not onto D2-MSNs (Figures 3 & 4).

First, we identified a significant increase in the amplitude of sEPSCs in D1-MSNs of mice treated with interrupted morphine, compared to mice treated with continuous morphine (Figure 3C). In contrast to this divergent adaptation, a convergent decrease in the AMPAR/NMDAR ratio was observed in D1-MSNs of male mice after both continuous and interrupted morphine treatment (Figure 4B). Since the patterns of morphine administration were divergent at the level of sEPSC amplitude, but convergent when measuring the AMPAR/NMDAR ratio, our findings suggest potential changes in the number and/or function of NMDARs. However, we did not detect any changes in the decay kinetics of the NMDAR EPSC that would indicate altered subunit composition (Table 3). Interestingly, our results differ from a previous report that identified an increased AMPAR/NMDAR ratio in D2- but not D1-MSNs at early withdrawal from repeated morphine administration (Zhu et al., 2016). Whilst this could simply reflect a difference in the pattern of administration, Zhu et al. (2016) found this adaptation was only observed at excitatory synaptic inputs from the paraventricular nucleus of the thalamus. This level of specificity could have been obscured in our study, which used electrical stimulation to activate all synapses concurrently. Future studies could use optogenetic stimulation to investigate the input-specificity of these cell type-specific adaptations evoked by continuous and interrupted morphine exposure.

Another adaptation conserved between both patterns of morphine administration, as well as both sexes, was a decrease in the paired-pulse ratio for D1-MSNs (Figure 4E). This indication of augmented glutamate release probability onto D1-MSNs has also been observed following extended withdrawal from repeated intermittent morphine injections (Hearing et al., 2016). Our findings suggest that these presynaptic adaptations occur in a consistent manner across different patterns of morphine administration (continuous, interrupted or intermittent) and different phases of withdrawal. Surprisingly, we did not identify a concurrent increase in the frequency of sEPSCs or the coefficient of variation for AMPAR responses, but these measures are both influenced by both the probability of glutamate release and the number of synaptic contacts. Opioid exposure can reduce dendritic spine density on NAc MSNs (Robinson and Kolb, 1999; Robinson et al., 2002; Graziane et al., 2016), and this decrease in the number of synaptic contacts may mask increases in sEPSC frequency related to the probability of glutamate release. We did not observe any changes in the rectification of AMPAR currents that would indicate synaptic incorporation of receptors lacking the GluA2 subunit (Table 2), but changes in AMPAR subunit composition may contribute to morphine-evoked plasticity in other phases of opioid administration and withdrawal (Hearing et al., 2016; Russell et al., 2016).

### Excitatory/inhibitory balance

NAc MSN functional states are regulated by the finely tuned integration of excitatory and inhibitory synaptic inputs. Disruptions in this excitatory/inhibitory balance can shift the functional states of MSNs and have been observed after psychostimulant exposure (Otaka et al., 2013; Yu et al., 2017). Here we showed that after six days of continuous morphine exposure, the NAc D1-MSN excitatory/inhibitory ratio was significantly decreased in male mice and increased in female mice (Figure 6A). This sex difference was surprising given both sexes showed convergent increases in presynaptic glutamate release (Figure 4E) and sIPSC amplitude (Figure 5C). This would suggest substantive sex-dependent adaptations in either post-synaptic AMPARs or GABA pre-synaptic release probability. Though repeated morphine exposure has been shown to induce transient silent excitatory synapses at early withdrawal, this effect is limited to D2- and not D1-MSNs and is therefore not a likely explanation for the reduced E/I balance observed in this study (Graziane et al., 2016). Together, the decrease in AMPAR/NMDAR and AMPAR/GABAR ratio would suggest that, in male mice, continuous morphine administration weakens the functional output of D1-MSNs, and potentially impairs reward-related behaviors linked to D1-MSN activity. Interrupted morphine administration did not induce an overall change in excitatory/inhibitory ratio in either MSN subtype. Given that interrupted morphine decreased the AMPAR/NMDAR but not AMPAR/GABAR ratio, it would appear that interrupted morphine does not affect excitatory input to D1-MSNs in the same fashion as continuous morphine. These differences in synaptic excitation and inhibition of D1-MSNs after continuous versus interrupted morphine exposure may contribute to the tolerance and sensitization of opioid reward that have been respectively reported after each pattern of exposure (Shippenberg et al., 1988; Lett, 1989; Gaiardi et al., 1991; Shippenberg et al., 1996; Russo et al., 2007; Sun et al., 2014; Yu et al., 2014).

### Sex Differences

An important feature of our experimental design was the inclusion of both female and male mice throughout all analyses. We noted some sex differences in basal synaptic transmission that were independent of morphine treatment: males had higher sIPSC frequency in D1-MSNs and sEPSC frequency in D2-MSNs, compared to their female counterparts. We also found evidence supporting sex-dependent effects of morphine exposure on synaptic plasticity: the AMPAR/GABAR ratio and AMPAR/NMDAR ratio were only decreased after continuous morphine exposure in male mice. Since both sexes developed a similar degree of psychomotor tolerance after continuous morphine exposure, the cellular changes that mediate psychomotor tolerance are clearly distinct, and may involve other circuits in female mice. Consistent with prior investigations of opioid-evoked synaptic plasticity, our study utilized mice at a peri-adolescent age, but it should be noted that the opioid-evoked adaptations observed here in adolescent mice could differ from those in older adult mice (Mayer-Blackwell et al., 2013, Zhang et al., 2009). Given that both clinical and preclinical studies report sex differences in opioid use disorders and addiction-like behaviors, it is critical that future studies continue to explore opioid evoked synaptic plasticity in both male and female rodents (Nicolas et al., 2022).

### Conclusions

In addition to psychomotor activation, mesolimbic dopamine signaling and differential gene expression (Lefevre et al., 2020), our present study shows that opioid-evoked synaptic plasticity is modulated by the pattern of morphine administration. Our data suggest that continuous morphine administration evokes adaptations that dampen D1-MSN functional output, and therefore could reduce subsequent reward-related behaviors. These adaptations in D1-MSN function are reduced by the interruption of continuous morphine exposure, which may enhance responses to subsequent opioid administration. Overall this study supports the hypothesis that maintaining continuity of opioid administration could be an effective therapeutic strategy to minimize the vulnerability to opioid use disorders.

## Conflict of interest statement

The authors declare no competing financial interests.

## Acknowledgements

Research reported in this publication was supported by the University of Minnesota’s MnDRIVE (Minnesota’s Discovery, Research, and Innovation Economy) initiative, as well as National Institutes of Health grants K99 DA052624 (EML), R00 DA037279 (PER,) and R01 DA048946 (PER). We thank David Leipold and Kerry Trotter for technical assistance.

## Notes

### Competing Interest Statement

The authors have declared no competing interest.

## REFERENCES

Ackerman SJ, Mordin M, Reblando J, Xu X, Schein J, Vallow S, Brennan M (2003) Patient-reported utilization patterns of fentanyl transdermal system and oxycodone hydrochloride controlled-release among patients with chronic nonmalignant pain. J Manag Care Pharm 9:223–231.

Burke DA, Rotstein HG, Alvarez VA (2017) Striatal Local Circuitry: A New Framework for Lateral Inhibition. Neuron 96:267–284.

Cahill CM (2020) Opioid dose regimen shapes mesolimbic adaptations. Neuropsychopharmacology 45:1777–1778.

Charbogne P et al. (2017) Mu Opioid Receptors in Gamma-Aminobutyric Acidergic Forebrain Neurons Moderate Motivation for Heroin and Palatable Food. Biol Psychiatry 81:778–788.

Cole SL, Robinson MJF, Berridge KC (2018) Optogenetic self-stimulation in the nucleus accumbens: D1 reward versus D2 ambivalence. PLoS One 13:e0207694.

Contet C, Filliol D, Matifas A, Kieffer BL (2008) Morphine-induced analgesic tolerance, locomotor sensitization and physical dependence do not require modification of mu opioid receptor, cdk5 and adenylate cyclase activity. Neuropharmacology 54:475–486.

Dong Y, Green T, Saal D, Marie H, Neve R, Nestler EJ, Malenka RC (2006) CREB modulates excitability of nucleus accumbens neurons. Nat Neurosci 9:475–477.

Enoksson T, Bertran-Gonzalez J, Christie MJ (2012) Nucleus accumbens D2- and D1-receptor expressing medium spiny neurons are selectively activated by morphine withdrawal and acute morphine, respectively. Neuropharmacology 62:2463–2471.

Evans CJ, Cahill CM (2016) Neurobiology of opioid dependence in creating addiction vulnerability. F1000Res 5.

Gaiardi M, Bartoletti M, Bacchi A, Gubellini C, Costa M, Babbini M (1991) Role of repeated exposure to morphine in determining its affective properties: place and taste conditioning studies in rats. Psychopharmacology (Berl) 103:183–186.

Gong S, Zheng C, Doughty ML, Losos K, Didkovsky N, Schambra UB, Nowak NJ, Joyner A, Leblanc G, Hatten ME, Heintz N (2003) A gene expression atlas of the central nervous system based on bacterial artificial chromosomes. Nature 425:917–925.

Graziane NM, Sun S, Wright WJ, Jang D, Liu Z, Huang YH, Nestler EJ, Wang YT, Schluter OM, Dong Y (2016) Opposing mechanisms mediate morphine- and cocaine-induced generation of silent synapses. Nat Neurosci 19:915–925.

Hearing MC, Jedynak J, Ebner SR, Ingebretson A, Asp AJ, Fischer RA, Schmidt C, Larson EB, Thomas MJ (2016) Reversal of morphine-induced cell-type-specific synaptic plasticity in the nucleus accumbens shell blocks reinstatement. Proc Natl Acad Sci U S A 113:757–762.

Heng LJ, Yang J, Liu YH, Wang WT, Hu SJ, Gao GD (2008) Repeated morphine exposure decreased the nucleus accumbens excitability during short-term withdrawal. Synapse 62:775–782.

Hikida T, Kimura K, Wada N, Funabiki K, Nakanishi S (2010) Distinct roles of synaptic transmission in direct and indirect striatal pathways to reward and aversive behavior. Neuron 66:896–907.

Kibaly C, Alderete JA, Liu SH, Nasef HS, Law PY, Evans CJ, Cahill CM (2021) Oxycodone in the Opioid Epidemic: High ‘Liking’, ‘Wanting’, and Abuse Liability. Cell Mol Neurobiol 41:899–926.

Koo JW, Lobo MK, Chaudhury D, Labonte B, Friedman A, Heller E, Pena CJ, Han MH, Nestler EJ (2014) Loss of BDNF signaling in D1R-expressing NAc neurons enhances morphine reward by reducing GABA inhibition. Neuropsychopharmacology 39:2646–2653.

Koob GF (2020) Neurobiology of Opioid Addiction: Opponent Process, Hyperkatifeia, and Negative Reinforcement. Biol Psychiatry 87:44–53.

Kourrich S, Thomas MJ (2009) Similar neurons, opposite adaptations: psychostimulant experience differentially alters firing properties in accumbens core versus shell. J Neurosci 29:12275–12283.

Kramer PF, Christensen CH, Hazelwood LA, Dobi A, Bock R, Sibley DR, Mateo Y, Alvarez VA (2011) Dopamine D2 receptor overexpression alters behavior and physiology in Drd2-EGFP mice. J Neurosci 31:126–132.

Le Marec T, Marie-Claire C, Noble F, Marie N (2011) Chronic and intermittent morphine treatment differently regulates opioid and dopamine systems: a role in locomotor sensitization. Psychopharmacology (Berl) 216:297–303.

Le Moine C, Bloch B (1995) D1 and D2 dopamine receptor gene expression in the rat striatum: sensitive cRNA probes demonstrate prominent segregation of D1 and D2 mRNAs in distinct neuronal populations of the dorsal and ventral striatum. J Comp Neurol 355:418–426.

Lefevre EM, Pisansky MT, Toddes C, Baruffaldi F, Pravetoni M, Tian L, Kono TJY, Rothwell PE (2020) Interruption of continuous opioid exposure exacerbates drug-evoked adaptations in the mesolimbic dopamine system. Neuropsychopharmacology 45:1781–1792.

Lett BT (1989) Repeated exposures intensify rather than diminish the rewarding effects of amphetamine, morphine, and cocaine. Psychopharmacology (Berl) 98:357–362.

Lichtblau L, Sparber SB (1981) Opiate withdrawal in utero increases neonatal morbidity in the rat. Science 212:943–945.

Lobo MK, Nestler EJ (2011) The striatal balancing act in drug addiction: distinct roles of direct and indirect pathway medium spiny neurons. Front Neuroanat 5:41.

Luscher C, Malenka RC (2011) Drug-evoked synaptic plasticity in addiction: from molecular changes to circuit remodeling. Neuron 69:650–663.

Ma YY, Cepeda C, Chatta P, Franklin L, Evans CJ, Levine MS (2012) Regional and cell-type-specific effects of DAMGO on striatal D1 and D2 dopamine receptor-expressing medium-sized spiny neurons. ASN Neuro 4.

Madayag AC, Gomez D, Anderson EM, Ingebretson AE, Thomas MJ, Hearing MC (2019) Cell-type and region-specific nucleus accumbens AMPAR plasticity associated with morphine reward, reinstatement, and spontaneous withdrawal. Brain Struct Funct 224:2311–2324.

Mayer-Blackwell B, Schlussman SD, Butelman ER, Ho A, Ott J, Kreek MJ, Zhang Y (2014) Self administration of oxycodone by adolescent and adult mice affects striatal neurotransmitter receptor gene expression. Neuroscience 258:280–291.

McDevitt DS, Jonik B, Graziane NM (2019) Morphine Differentially Alters the Synaptic and Intrinsic Properties of D1R- and D2R-Expressing Medium Spiny Neurons in the Nucleus Accumbens. Front Synaptic Neurosci 11:35.

Mu P, Moyer JT, Ishikawa M, Zhang Y, Panksepp J, Sorg BA, Schluter OM, Dong Y (2010) Exposure to cocaine dynamically regulates the intrinsic membrane excitability of nucleus accumbens neurons. J Neurosci 30:3689–3699.

Nelson AB, Hang GB, Grueter BA, Pascoli V, Luscher C, Malenka RC, Kreitzer AC (2012) A comparison of striatal-dependent behaviors in wild-type and hemizygous Drd1a and Drd2 BAC transgenic mice. J Neurosci 32:9119–9123.

Nicolas C, Zlebnik NE, Farokhnia M, Leggio L, Ikemoto S, Shaham Y (2022) Sex Differences in Opioid and Psychostimulant Craving and Relapse: A Critical Review. Pharmacol Rev 74:119–140.

O’Donnell P, Greene J, Pabello N, Lewis BL, Grace AA (1999) Modulation of cell firing in the nucleus accumbens. Ann N Y Acad Sci 877:157–175.

O’Neal TJ, Nooney MN, Thien K, Ferguson SM (2020) Chemogenetic modulation of accumbens direct or indirect pathways bidirectionally alters reinstatement of heroin-seeking in high-but not low-risk rats. Neuropsychopharmacology 45:1251–1262.

O’Neal TJ, Bernstein MX, MacDougall DJ, Ferguson SM (2022) A Conditioned Place Preference for Heroin Is Signaled by Increased Dopamine and Direct Pathway Activity and Decreased Indirect Pathway Activity in the Nucleus Accumbens. J Neurosci 42:2011–2024.

Otaka M, Ishikawa M, Lee BR, Liu L, Neumann PA, Cui R, Huang YH, Schluter OM, Dong Y (2013) Exposure to cocaine regulates inhibitory synaptic transmission in the nucleus accumbens. J Neurosci 33:6753–6758.

Pisansky MT, Lefevre EM, Retzlaff CL, Trieu BH, Leipold DW, Rothwell PE (2019) Nucleus Accumbens Fast-Spiking Interneurons Constrain Impulsive Action. Biol Psychiatry 86:836–847.

Robinson TE, Kolb B (1999) Morphine alters the structure of neurons in the nucleus accumbens and neocortex of rats. Synapse 33:160–162.

Robinson TE, Gorny G, Savage VR, Kolb B (2002) Widespread but regionally specific effects of experimenter-versus self-administered morphine on dendritic spines in the nucleus accumbens, hippocampus, and neocortex of adult rats. Synapse 46:271–279.

Rothwell PE, Gewirtz JC, Thomas MJ (2010) Episodic withdrawal promotes psychomotor sensitization to morphine. Neuropsychopharmacology 35:2579–2589.

Rumbaugh G, Vicini S (1999). Distinct synaptic and extrasynaptic NMDA receptors in developing cerebellar granule neurons. J Neurosci 19:10603–10610.

Russell SE, Puttick DJ, Sawyer AM, Potter DN, Mague S, Carlezon WA, Jr., Chartoff EH (2016) Nucleus Accumbens AMPA Receptors Are Necessary for Morphine-Withdrawal-Induced Negative-Affective States in Rats. J Neurosci 36:5748–5762.

Russo SJ, Bolanos CA, Theobald DE, DeCarolis NA, Renthal W, Kumar A, Winstanley CA, Renthal NE, Wiley MD, Self DW, Russell DS, Neve RL, Eisch AJ, Nestler EJ (2007) IRS2-Akt pathway in midbrain dopamine neurons regulates behavioral and cellular responses to opiates. Nat Neurosci 10:93–99.

Sesack SR, Grace AA (2010) Cortico-Basal Ganglia reward network: microcircuitry. Neuropsychopharmacology 35:27–47.

Shippenberg TS, Heidbreder C, Lefevour A (1996) Sensitization to the conditioned rewarding effects of morphine: pharmacology and temporal characteristics. Eur J Pharmacol 299:33–39.

Shippenberg TS, Emmett-Oglesby MW, Ayesta FJ, Herz A (1988) Tolerance and selective cross-tolerance to the motivational effects of opioids. Psychopharmacology (Berl) 96:110–115.

Shuen JA, Chen M, Gloss B, Calakos N (2008) Drd1a-tdTomato BAC transgenic mice for simultaneous visualization of medium spiny neurons in the direct and indirect pathways of the basal ganglia. J Neurosci 28:2681–2685.

Soares-Cunha C, Coimbra B, David-Pereira A, Borges S, Pinto L, Costa P, Sousa N, Rodrigues AJ (2016) Activation of D2 dopamine receptor-expressing neurons in the nucleus accumbens increases motivation. Nat Commun 7:11829.

Soares-Cunha C, de Vasconcelos NAP, Coimbra B, Domingues AV, Silva JM, Loureiro-Campos E, Gaspar R, Sotiropoulos I, Sousa N, Rodrigues AJ (2020) Nucleus accumbens medium spiny neurons subtypes signal both reward and aversion. Mol Psychiatry 25:3241–3255.

Sun L, Hu L, Li Y, Cui C (2014) Mesoaccumbens dopamine signaling alteration underlies behavioral transition from tolerance to sensitization to morphine rewarding properties during morphine withdrawal. Brain Struct Funct 219:1755–1771.

Tai LH, Lee AM, Benavidez N, Bonci A, Wilbrecht L (2012) Transient stimulation of distinct subpopulations of striatal neurons mimics changes in action value. Nat Neurosci 15:1281–1289.

Toddes C, Lefevre EM, Brandner DD, Zugschwert L, Rothwell PE (2021) mu-Opioid Receptor (Oprm1) Copy Number Influences Nucleus Accumbens Microcircuitry and Reciprocal Social Behaviors. J Neurosci 41:7965–7977.

Trieu BH, Remmers BC, Toddes C, Brandner DD, Lefevre EM, Kocharian A, Retzlaff CL, Dick RM, Mashal MA, Gauthier EA, Xie W, Zhang Y, More SS, Rothwell PE (2022) Angiotensin-converting enzyme gates brain circuit-specific plasticity via an endogenous opioid. Science 375:1177–1182.

Vanderschuren LJ, Tjon GH, Nestby P, Mulder AH, Schoffelmeer AN, De Vries TJ (1997) Morphine-induced long-term sensitization to the locomotor effects of morphine and amphetamine depends on the temporal pattern of the pretreatment regimen. Psychopharmacology (Berl) 131:115–122.

Wang J, Ishikawa M, Yang Y, Otaka M, Kim JY, Gardner GR, Stefanik MT, Milovanovic M, Huang YH, Hell JW, Wolf ME, Schluter OM, Dong Y (2018) Cascades of Homeostatic Dysregulation Promote Incubation of Cocaine Craving. J Neurosci 38:4316–4328.

Wolf ME, Tseng KY (2012) Calcium-permeable AMPA receptors in the VTA and nucleus accumbens after cocaine exposure: when, how, and why? Front Mol Neurosci 5:72.

Yu G, Zhang FQ, Tang SE, Lai MJ, Su RB, Gong ZH (2014) Continuous infusion versus intermittent bolus dosing of morphine: a comparison of analgesia, tolerance, and subsequent voluntary morphine intake. J Psychiatr Res 59:161–166.

Yu J, Yan Y, Li KL, Wang Y, Huang YH, Urban NN, Nestler EJ, Schluter OM, Dong Y (2017) Nucleus accumbens feedforward inhibition circuit promotes cocaine self-administration. Proc Natl Acad Sci U S A 114:E8750–E8759.

Yuan T, Bellone C (2013) Glutamatergic receptors at developing synapses: the role of GluN3A-containing NMDA receptors and GluA2-lacking AMPA receptors. Eur J Pharmacol 719:107–111.

Zhang XF, Hu XT, White FJ (1998) Whole-cell plasticity in cocaine withdrawal: reduced sodium currents in nucleus accumbens neurons. J Neurosci 18:488–498.

Zhang Y, Picetti R, Butelman ER, Schlussman SD, Ho A, Kreek MJ (2009) Behavioral and neurochemical changes induced by oxycodone differ between adolescent and adult mice. Neuropsychopharmacology 34:912–922.

Zhu Y, Wienecke CF, Nachtrab G, Chen X (2016) A thalamic input to the nucleus accumbens mediates opiate dependence. Nature 530:219–222.

